# Exploring the Role of Hypusine Signaling in Vascular Smooth Muscle Cells for Mitigating Restenosis in Coronary Artery Disease

**DOI:** 10.64898/2025.12.15.694523

**Authors:** Yann Grobs, Sarah-Eve Lemay, Manon Mougin, Charlie Theberge, Magalie Boucher, Sandra Breuils-Bonnet, Sandra Martineau, Alice Bourgeois, Andreanne Pelletier, Maud Fillon, Jean Perron, François Potus, Steeve Provencher, Olivier Boucherat, Sebastien Bonnet

## Abstract

**Background:** Post-surgical restenosis in patients with coronary artery disease (CAD) is a pathological vascular remodeling process characterized by neointimal hyperplasia, mainly driven by vascular smooth muscle cells (VSMCs) phenotypic switching toward synthetic and proliferative state. This study identifies novel signaling pathway promoting pro-proliferative phenotype of VSMC and contributing to the neointimal hyperplasia development.

**Methods:** The expression of hypusine signaling components was evaluated in human primary culture of coronary artery smooth muscle cells (CoASMCs) isolated from controls and patients with CAD, using comparative proteomic analysis and western blotting, as well as in three preclinical animal models of restenosis; rat carotid injury, mice carotid ligation and canine coronary artery bypass graft. CAD-CoASMCs proliferation was assessed by western blot and immunofluorescence with pharmacological (GC7) and molecular (shRNA) inhibitors of deoxyhypusine synthase (DHPS). The contribution of hypusine signaling to neointimal hyperplasia was investigated using both pharmacological and smooth muscle cell-specific knockout mice approaches. Additionally, human saphenous vein and human coronary artery tissue cultures were employed to explore the translational potential of targeting hypusine signaling to prevent neointimal hyperplasia.

**Results:** All components of the hypusine pathway (eukaryote translational initiation factor 5A (eIF5A), deoxyhypusine hydroxylase (DOHH) and DHPS) were significantly overexpressed in CAD-CoASMCs and in preclinical animal models of restenosis. Pharmacological and molecular inhibition of DHPS reduced eIF5A hypusination, VSMC proliferation and expression of extracellular matrix proteins. Proteomic and KEGG analyses demonstrated disruption of cell cycle and DNA replication pathways, including a downregulation of threonine tyrosine kinase (TTK). Our findings suggest that TTK acts as a downstream effector of hypusine signaling, partly mediating to the proliferative effects observed in CAD-CoASMCs. *In vivo*, pharmacological and genetic inhibition of DHPS significantly reduced neointimal hyperplasia without adverse effects. Finally, *ex vivo* human tissue culture confirmed that GC7 mitigates growth factor–induced vascular remodeling.

**Conclusions:** Hypusine signaling is a critical regulator of VSMC proliferation for neointimal hyperplasia. Inhibiting DHPS reduces vascular remodeling, making it a promising target for preventing restenosis after coronary interventions.

**Clinical Perspective:** *What Is New?:*

- Hypusine signaling is markedly upregulated in coronary artery smooth muscle cells (CoASMCs) from patients with coronary artery disease (CAD) and in multiple preclinical models of restenosis.
- Proteomic profiling identifies DHPS, the rate-limiting enzyme for eIF5A hypusination, as a key driver of vascular smooth muscle cell (VSMC) pro-proliferative phenotype and extracellular matrix production.
- Pharmacological (GC7) and genetic inhibition of DHPS effectively suppress eIF5A hypusination, attenuate the synthetic and proliferative CAD-CoASMCs phenotype, and significantly reduce neointimal hyperplasia in rodent models of vascular injury.
- *Ex vivo* human tissue demonstrates that DHPS inhibition prevents neointimal hyperplasia, providing strong translational evidence.

*What Are the Clinical Implications?:*

- These findings establish hypusine signaling as a previously unrecognized regulator of pathological VSMC activation in CAD and restenosis.
- DHPS inhibition emerges as a promising therapeutic strategy to prevent neointimal hyperplasia following coronary interventions such as angioplasty, stenting, or bypass grafting.
- Collectively, our data support the clinical development of selective DHPS inhibitors as a novel class of therapeutics to improve long-term outcomes after coronary revascularization and potentially other occlusive vascular diseases.

## INTRODUCTION

Coronary artery disease (CAD) is the predominant form of cardiovascular disease and remains the principal cause of mortality worldwide^1^. For patients with severe coronary stenosis, revascularization procedures including balloon angioplasty, stent implantation, and coronary artery bypass grafting (CABG) are often lifesaving. However, these interventions are frequently compromised by restenosis, a pathological re-narrowing of the vessel driven by exuberant neointimal hyperplasia and associated with recurrent ischemic events such as myocardial infarction^2–5^. Although drug-eluting stents have significantly reduced early restenosis rates, excessive vascular smooth muscle cell (VSMC) proliferation coupled with impaired re-endothelialisation continues to hinder long-term clinical outcomes^3^. These limitations underscore the need for therapies that target the molecular drivers underlying restenosis.

Under physiological conditions, VSMCs maintain a highly specialized contractile phenotype essential for vascular tone and homeostasis^6^. however, following vascular injury like after stent implantation, VSMCs undergo a profound phenotypic switch toward a synthetic, repair-oriented state^7^. This transition is characterized by increased proliferation, cell migration, extracellular matrix (ECM) deposition, metabolic reprogramming toward glycolysis, enhanced DNA repair^6–13^. Sustained activation of this maladaptive phenotype is a core driver of neointimal hyperplasia in both atherosclerosis and restenosis^7,8,14–16^ and contributes to several vascular diseases including aortic aneurysm^17^. Yet a unifying molecular mechanism capable of integrating these proliferative, metabolic, and contractile abnormalities in CAD-associated SMCs remains undefined. While transcriptomic shifts in VSMC pro-proliferative phenotype are well described, the translational control layer ultimately determining protein output remains largely unexplored.

In this present study, we performed comparative proteomic analysis to identify novel targets that drive the proliferative state of VSMCs contributing to neointimal hyperplasia in restenosis. Interestingly, pathological state of CAD-CoASMCs was accompanied by significant upregulation of deoxyhypusine synthase (DHPS) and increased eukaryotic translation initiation factor 5A (eIF5A) hypusination, a signaling pathway recently highlighted as a therapeutic target in cancer biology and pulmonary vascular disease. These two pathologies share striking features with CAD-derived CoASMCs, including hyperproliferation and metabolic impairment pointing to a central role for this post-translational modification of eIF5A^9,18^. Hypusination involves the formation of a unique amino acid residue on eIF5A through sequential enzymatic steps catalyzed by DHPS and deoxyhypusine hydroxylase (DOHH). First, DHPS converts eIF5A in deoxyhypusine-eIF5A, and subsequently, DOHH catalyzes the formation of hypusinated eIF5A (Figure S1). This modification is critical for the activation and cytoplasmic localization of eIF5A and is dependent on spermidine, a polyamine synthesized within the polyamine metabolic pathway, known to promote proliferation of VSMC in restenosis^19–22^ (Figure S1). Once hypusinated, eIF5A selectively enhances the translation of polyproline-rich proteins involved in cell-cycle progression, proliferation, and cytoskeletal remodeling^18,23–25^. The inhibition of DHPS or DOHH impairs eIF5A activation and strongly suppresses proliferation, highlighting this pathway as an attractive therapeutic target^18,25^. Through complementary pharmacological and molecular inhibition of DHPS, we identified hypusine signaling as a central regulator of CAD-CoASMC proliferation. Importantly, inactivation of eIF5A conferred substantial protection against neointimal hyperplasia in two preclinical models of restenosis and in *ex vivo* cultured human coronary artery (CoA) and saphenous vein (SV) tissues.

Collectively, these findings reveal the hypusination axis as a previously unrecognized determinant of VSMC behavior and post-intervention pathological vascular remodeling. Targeting this translational control mechanism may provide a novel therapeutic strategy to prevent restenosis and improve long-term outcomes in patients with CAD.

## METHODS

### Human Tissue

Human samples, including saphenous veins were collected from patients undergoing CABG surgery and coronary arteries were obtained from explanted hearts during transplantation. The diagnosis of CAD was validated through clinical history, angiography, and criteria such as previous stent placement or bypass grafts, confirmed by experienced cardiologists. Experimental procedures involving human tissues or cells adhered to the guidelines established by the Declaration of Helsinki and received approval from the Biosafety and Ethics Committees of Laval University and the Institut Universitaire de Cardiologie et de Pneumologie de Québec (approval numbers CER22376 and CER20841) with signed informed consent was obtained. Details about patient characteristics are provided in Table S3.

### Animal models

The sample size for each group was determined to identify at least a 20% difference between experimental and control conditions, with 80% statistical power and a standard deviation of 10%. Animals of the same sex and genotype, with comparable body weights, were generated and randomly assigned to various experimental groups. No animals were excluded from the analyses. Investigators conducting histological assessments were blinded to the group assignments to prevent bias. All animal experiments received approval from the Animal Ethics Committee at Université Laval (approval numbers #2020-616) and were conducted in accordance with the guidelines established by the Canadian Council on Animal Care and the ARRIVE (Animal Research: Reporting of In Vivo Experiments) guidelines.

### Statistical analysis

The normality of the data distribution was evaluated using the Shapiro-Wilk test. When comparing the means of two groups with normally distributed data, an unpaired Student’s t-test was employed. For comparisons across multiple groups, one-way ANOVA was conducted, followed by Tukey’s post hoc tests. Pearson correlation analysis was employed to assess relationships between variables. Results are expressed as mean ± SEM. A p-value of less than 0.05 was considered statistically significant. Significance levels are denoted as *P<0.05, **P<0.01, ***P<0.001, and ****P<0.0001. All graphical representations and statistical analyses were performed using GraphPad Prism version 10.6.1.

In the case of paired data (Figure 7 and 8), data were reported using means and standard deviations or medians and interquartile ranges according to the variable distributions. The three conditions for each subject were analyzed using a linear mixed model with one fixed factor for the comparison among conditions and one random effect for variability among subjects. The dependence among residuals of repeated measurements within the same experimental unit were estimated with an unstructured covariance association. The normality hypothesis was verified using the Shapiro-Wilks test using residuals from the statistical model and transformed by the Cholesky’s metric. The Brown and Forsythe’s variation of Levene’s test statistic was used to verify the homogeneity of variances. Statistical significance was declared significant with a two-tailed *p* value < 0.05. Analyses were performed using SAS version 9.4 (SAS Institute Inc, Cary, NC, U.S.A.).

## RESULTS

### The Hyper-Proliferative Phenotype of CAD-CoASMCs Is Associated with Enhanced Hypusine Signaling

As a starting point, we characterized the pathological phenotype of our CoASMCs isolated from patients with CAD. Compared to CTRL-CoASMCs, CAD-CoASMCs exhibited reduced expression of contractile markers as shown by western blot (α-smooth muscle actin (α-SMA), and calponin), and RT-qPCR (myocardin; *MYOCD*)^7,10–12,14^ (Figure 1A and Figure S2). Conversely, markers of proliferation such as minichromosome maintenance complex component 2 (MCM2) and polo-like kinase 1 (PLK1), proliferating cell nuclear antigen (PCNA), and survivin (*BIRC5*) were increased (Figure 1A and Figure S2).

**Figure 1.**
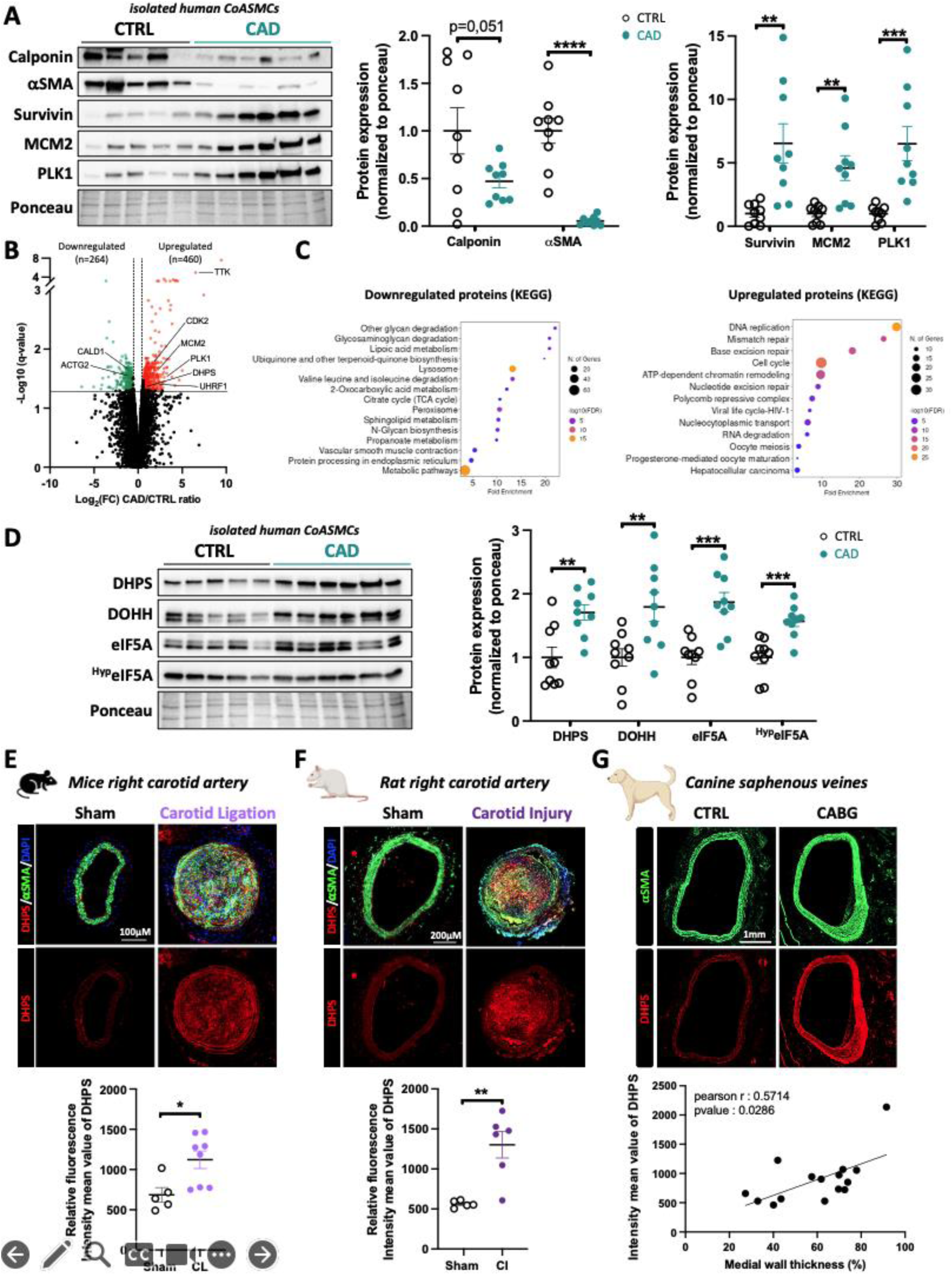
Overexpressed hypusine signaling in coronary artery smooth muscle cells of CAD patients and animal models. (**A**) Representative western blot and corresponding quantification of calponin, αSMA, survivin, MCM2, PLK1 in isolated CoASMCs from control and CAD patients (*n=9; **P < 0.01, ***P < 0.001, ****P < 0.0001, unpaired Student’s t-test; data represent mean ± SEM)*. (**B**) Volcano plot of the *q*-values versus log_2_ fold change (FC) protein abundance differences between CTRL-CoASMCs (n=4) and CAD-CoASMCs (n=5), with proteins considered as differentially expressed (fold change > 1.5, *q-*value < 0,05) indicated in green (downregulated) or red (upregulated). Proteins of interest are identified. (**C**) Enrichment analysis performed by Shinigo software v0.85 showing the top of KEGG pathways in both down- and upregulated proteins. KEGG pathways are ranked by fold enrichment values. The most significant processes are highlighted in orange, and the less significant processes are highlighted in purple according to -log_10_ (false discovery rate) values. (**D**) Representative western blot and corresponding quantification of DHPS, DOHH, eIF5A and Hypusinated form of eIF5a (^Hyp^eIF5A in isolated CoASMCs, *(n = 9; *P < 0.05 and ***P < 0.001, unpaired Student’s t-test; data represent mean ± SEM)*. (**E**) Representative immunofluorescence images of DHPS (Red), αSMA (green) in carotid artery from mice subjected or not to carotid ligation. DAPI (4′,6-diamidino- 2-phenylindole) (blue) was used to detect nuclei *(n=5 to 8; *P < 0.05, unpaired Student’s t-test; data represent mean ± SEM)*. Scale bar, 100µm. (**F**) Representative immunofluorescence images of DHPS (Red), αSMA (green) in carotid artery from rat subjected or not to carotid injury. DAPI (blue) was used to detect nuclei *(n=5 to 6; **P < 0.01, unpaired Student’s t-test; data represent mean ± SEM)*. Scale bar, 200µm. (**G**) Representative immunofluorescence images of DHPS (Red), αSMA (green) in carotid artery from in saphenous vein from canine subjected or not to coronary artery bypass graft (CABG). DAPI (blue) was used to detect nuclei. Correlation between DHPS fluorescence intensity and medial wall thickness is shown in the bottom *(n=15, Pearson’s, r = 0.5714; P<0.05)*. Scale bar, 1mm.

To identify molecular pathways and key proteins contributing to the pathological phenotype of CAD-CoASMCs, we performed nanoscale liquid chromatography tandem mass spectrometry (LC-MS/MS) on four CTRL and five CAD human CoASMCs lines. Using differential expression thresholds (absolute fold change > 1.5; q-value < 0.05), we identified 724 differentially expressed proteins (DEPs), including 460 upregulated and 264 downregulated in CAD-CoASMCs. A volcano plot illustrating these DEPs is shown in Figure 1A. Consistent with VSMC phenotypic switching, contractile proteins such as CALD1 (Caldesmon) and smooth muscle actin gamma-2^15,26^ (ACTG2) were significantly downregulated, while proliferative drivers including MCM2 and PLK1 were markedly increased. Additional proteins controlling VSMC plasticity in vascular remodeling and atherosclerosis such as Ubiquitin-like containing PHD and RING finger domains 1 (UHRF1)^27^, Cyclin-dependent kinase 2 (CDK2)^28,29^ and threonine tyrosine kinase (TTK)^10^ were found significantly increased in CAD-CoASMCs, further supporting abnormal phenotype of CAD-CoASMCs. Accordingly, functional annotation using ShinyGO (v0.85) showed that downregulated proteins mapped to VSMC contraction, whereas upregulated proteins were enriched in DNA replication, cell cycle progression, and DNA repair pathways (Figure 1B). All these findings confirm that CAD-CoASMCs adopt a pro-proliferative and synthetic phenotype characteristic of neointimal hyperplasia, a key process of restenosis

Among the upregulated proteins, DHPS emerged as a prominent candidate due to its involvement in hypusine signaling recently highlighted as a therapeutic target in diseases characterized by a hyperproliferative state like cancer et pulmonary hypertension. The increase of hypusine signaling was confirmed by western blots showing significantly increased DHPS, DOHH, total eIF5A, and hypusinated eIF5A in CAD-CoASMCs compared to CTRL (Figure 1D). Immunofluorescence staining revealed increased DHPS and eIF5A expression in two animal models of restenosis: rat carotid endothelial denudation and mouse total carotid ligation (Figure 1E and F; Figure S3A and B). Elevated DHPS and eIF5A levels were also correlated with vascular remodeling in SV grafts in a canine CABG model (Figure 1G; Figure S3C).

Together these findings suggesting that the pathological phenotype seen in CAD-CoASMCs is associated with an upregulation of hypusine signaling.

### Hypusine signaling promotes proliferation of CAD-CoASMCs

Given the marked upregulation of hypusine signaling in CoASMCs isolated from patients with CAD and in animal models, we next investigated whether this pathway functionally contributes to the hyperproliferative phenotype of CAD-CoASMCs. We first used the well-established DHPS inhibitor N1-guanyl-1,7-diaminoheptane (GC7), a competitive spermidine analog^30^. As expected, exposure of CAD-CoASMCs to GC7 markedly suppressed eIF5A hypusination (^Hyp^eIF5A), while simultaneously increasing the inactive, acetylated form of eIF5A (^Ac^eIF5A), without altering total eIF5A protein levels (Figure 2A). Inhibition of hypusine signaling resulted in a significant decrease in the expression of genes involved in cell cycle progression such as MCM2, PLK1, PCNA, and survivin (Figure 2A). Immunofluorescence staining of Ki67 further confirmed that GC7 treatment significantly reduces CAD-CoASMC proliferation (Figure 2B).

**Figure 2.**
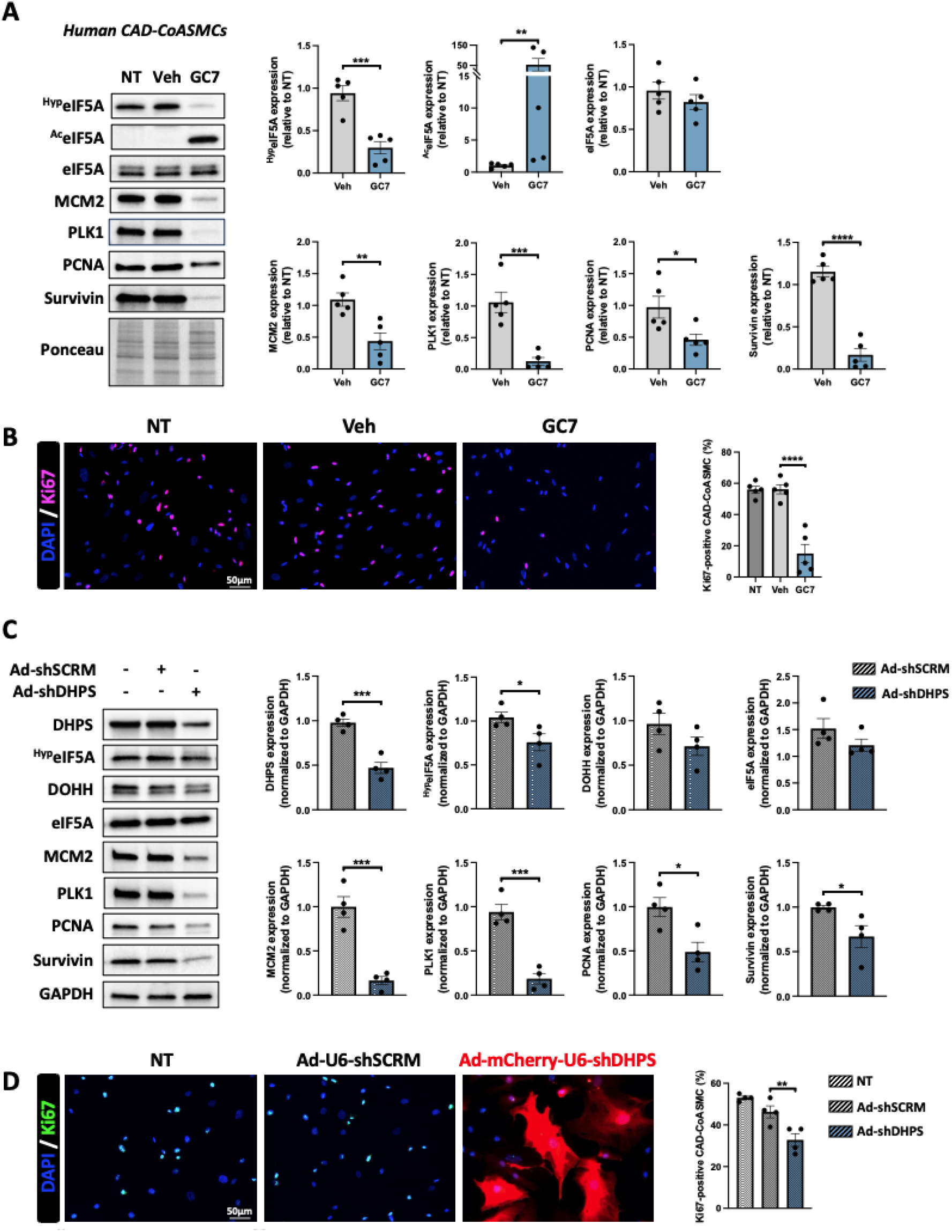
Hypusine signaling promotes CAD-CoASMCs proliferation. (**A**) Representative western blot and corresponding quantification of ^Hyp^eIF5A, ^Ac^eIF5A, eIF5A, MCM2, PLK1, PCNA and survivin in CAD-CoASMCs exposed or not to GC7 (30µM) for 48h. (*n=5; *P < 0.05, **P < 0.01, ***P < 0.001, ****P < 0.0001, unpaired Student’s t-test; data represent mean ± SEM)*. (**B**) Representative fluorescent images and corresponding quantification of Ki67- labeled (red) of CAD-CoASMCs exposed or not to GC7 (30µM) for 48h. Cell nuclei were counterstained with DAPI (blue) (*n = 5; ****P < 0.001, one-way ANOVA followed by Tukey’s post hoc analysis; data represent mean ± SEM*). Scale bar, 50μm. (**C**) Representative western blot and corresponding quantification of DHPS, ^Hyp^eIF5A, DOHH, eIF5A, MCM2, PLK1, PCNA, survivin and GAPDH in CAD-CoASMCs infected with an adenovirus encoding either a shRNA that targets DHPS (Ad-shDHPS) or a scrambled shRNA (Ad-shSCRM) for 96h. GAPDH served as a loading control. (*n=4; *P < 0.05, ***P < 0.001, unpaired Student’s t-test; data represent mean ± SEM)*. (**D**) Representative fluorescent images and corresponding quantification of Ki67- labeled (red) of CAD-CoASMCs infected with Ad-shDHPS or Ad-shSCRM for 96h. Cell nuclei were counterstained with DAPI (blue) (*n = 4; **P < 0.01, one-way ANOVA followed by Tukey’s post hoc analysis; data represent mean ± SEM*). Scale bar, 50μm.

Because phenotypic plasticity of VSMCs also involves increased ECM synthesis, we measured the protein expression levels of key ECM components in GC7-treated cells. As shown in Figure S3, collagen I (COL1), collagen III (COL3) and fibronectin (FN) were markedly decrease following GC7 exposure, indicating that activation of eIF5A contributes to both proliferative and matrix-synthetic programs in CAD-CoASMCs.

To validate that these effects were specifically due to DHPS inhibition, we performed molecular loss-of-function studies using an adenovirus encoding a DHPS-targeting shRNA. As anticipated, shDHPS effectively knocked down DHPS expression and reduced eIF5A hypusination (Figure 2C), without affecting expression of DOHH or total eIF5A, confirming target specificity. Consistent with pharmacological inhibition, DHPS knockdown decreased SMCs proliferation and lowered the expression of MCM2, PLK1, PCNA, and survivin (Figure 2C and 2D).

Together, these complementary approaches demonstrate that DHPS-dependent eIF5A hypusination is required to sustain the proliferative and ECM-producing phenotype of CAD-CoASMCs, positioning hypusine signaling as a central regulator of pathological VSMC activation.

### Pharmacological inhibition of DHPS using GC7 reshapes the proteome of CAD-CoASMCs

Because hypusinated eIF5A is required for the translation of a specific subset of mRNAs, we next conducted an analysis of the proteome changes in human CAD-CoASMCs induced in response to GC7. Using standard criteria for differential expression (absolute fold change > 1.5, q-value < 0.05), we identified 901 differentially expressed proteins (DEPs), of which 239 were upregulated and 662 downregulated following GC7 treatment (Figure S5A and B). Our proteomic dataset validated several findings obtained previously by Western blot. Proteins associated with proliferation, including MCM2 and PLK1, were markedly reduced in GC7-treated CAD-CoASMCs (Figure S5C), consistent with results shown in Figure 2A. Similarly, GC7 decreased expression of ECM-related proteins (COL1A1, COL1A2, COL3A1) and key proliferative regulators in CAD (CDK2, TTK), aligning with our earlier biochemical analyses. KEGG pathway analysis revealed that downregulated proteins were strongly enriched in pathways related to protein synthesis (ribosome) and cell metabolism, consistent with impaired translational capacity upon reduction of eIF5A hypusination. In contrast, upregulated proteins clustered in pathways related to oxidative stress responses and amino acid biosynthesis (Figure S5C). This pattern suggests that cells experiencing reduced hypusine-mediated translation activate compensatory metabolic programs to restore proteostasis. Supporting this interpretation, GC7-treated CAD-CoASMCs exhibited increased DHPS expression, likely reflecting a feedback mechanism attempting to counteract the effects of its inhibited activity.

To determine whether hypusine inhibition reverses the disease-associated proteomic signature, we compared DEPs from GC7-treated cells with proteins previously identified as upregulated in CAD-CoASMCs. Among 460 proteins elevated in CAD, 92 were downregulated by GC7 (Figure 3A and B). Pathway enrichment confirmed that these overlapping proteins predominantly regulate the cell cycle and DNA replication (Figure 3C), indicating that DHPS inhibition directly counteracts core drivers of the hyperproliferative phenotype. Notably, TTK, a dual-specificity kinase (serine, threonine, tyrosine) involved in several mechanisms of cell-cycle control^31–34^, was one of the most highly overexpressed proteins in CAD-CoASMCs and was also among the top 10 most significantly reduced proteins following GC7 exposure, suggesting a central role in mediating the antiproliferative effects of hypusine pathway inhibition.

**Figure 3.**
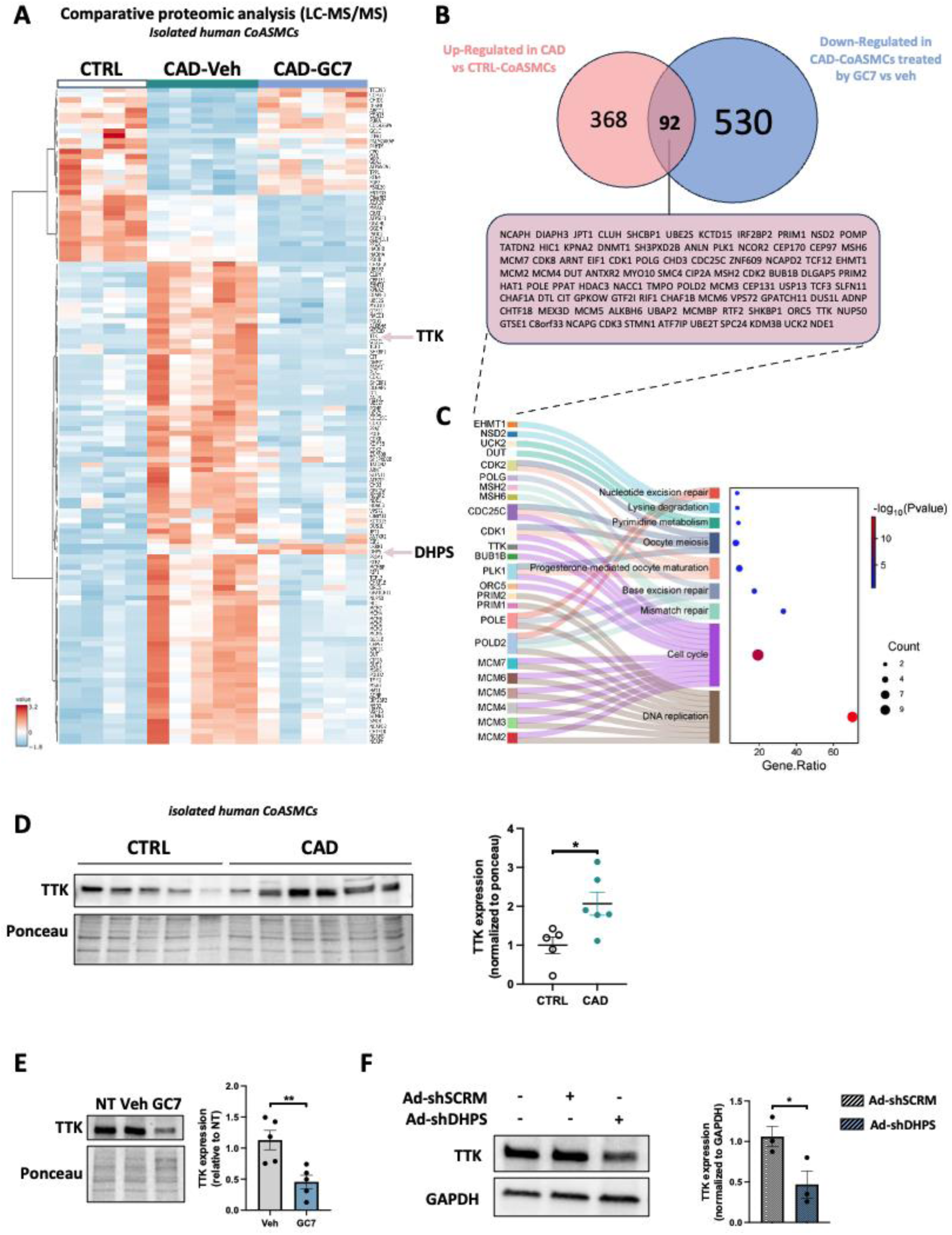
DHPS inhibition-mediated by GC7 reverses the proliferative protein expression signature in CAD-CoASMCs. **(A)** Heatmap depicting the differentially expressed proteins (log_2_ fold change > 1.5 and q-value < 0.05) as determined by nanoscale liquid chromatography coupled to tandem mass spectrometry (LC-MS/MS) analysis of CTRL-CoASMCs (n = 4) and CAD-CoASMCs (n = 5) subjected or not to GC7 (30µM) for 48h. Distance measurement method: Pearson; clustering method: average linkage. Dark blue to dark red color gradient denotes lower to higher expression. Proteins present in the heatmap are provided in the online supplement. **(B)** Venn diagram illustrating the overlap between proteins that are upregulated in CAD-CoASMCs compared to control (CTRL) and proteins that are downregulated in CAD-CoASMCs following 48h treatment with GC7 (30µM). The ninety-two common proteins are highlighted in the box. **(C)** Sankey dot plot performed with SRplot illustrates the association between differentially expressed proteins and enriched biological processes (KEGG pathway). Left part is sankey plot, represents proteins within each pathway. Right part is dot plot; dot sizes represent proteins numbers. The color coding indicates the biological functions. The position of dots represents the significance level of enrichment, to the right is the most significant. **(D)** Western blot and corresponding quantification of TTK in isolated CoASMCs from CTRL (n=5) and CAD patients (n=6) (*\*P < 0.05, unpaired Student’s t-test; data represent mean ± SEM)*. **(E)** Representative western blot and corresponding quantification of TTK in CAD-CoASMCs exposed or not to GC7 (30µM) for 48h. (*n=5, **P < 0.01, unpaired Student’s t-test; data represent mean ± SEM)*. **(F)** Representative western blot and corresponding quantification of TTK and GAPDH in CAD-CoASMCs infected with an adenovirus encoding either a shRNA that targets DHPS (Ad-shDHPS) or a scrambled shRNA (Ad-shSCRM) for 96h. GAPDH served as a loading control. (*n=3; *P < 0.05, unpaired Student’s t-test; data represent mean ± SEM)*.

TTK is typically undetectable in normal cells but is strongly overexpressed in proliferating cells, particularly in multiple cancers where it promotes tumor growth, metastasis, and poor prognosis^35–40^. Recent study has demonstrated that TTK regulates PLK1 activity, making it a compelling target in cancer therapy^41^. Addiotionally, TTK has also been implicated in neointimal hyperplasia and restenosis in mouse models, where it enhances VSMC proliferation^10^. Based on these observations, we hypothesized that the antiproliferative effects of DHPS inhibition in CAD-CoASMCs may be mediated, at least in part, by the downregulation of TTK. To test our hypothesis, we first validated by Western blot that TTK is significantly overexpressed in CAD-CoASMCs compared with CTRL-CoASMCs (Figure 3D). We then confirmed that GC7 markedly reduces TTK protein expression in CAD-CoASMCs (Figure 3E–F), mirroring our proteomic results.

To test the functional relevance of TTK suppression, we inhibited TTK using both the pharmacological inhibitor CFI-402257 and a TTK-specific siRNA (siTTK). Dose-response experiments with CFI-402257 showed a clear, dose-dependent reduction in TTK levels (Figure 4A), accompanied by decreases in MCM2, PLK1, and survivin. Molecular inhibition using siTTK efficiently reduced TTK expression (Figure 4B) and likewise lowered proliferation-associated proteins. Immunofluorescence analysis of Ki67 confirmed that both CFI-402257 and siTTK significantly impaired CAD-CoASMC proliferation (Figure 4C–D).

**Figure 4.**
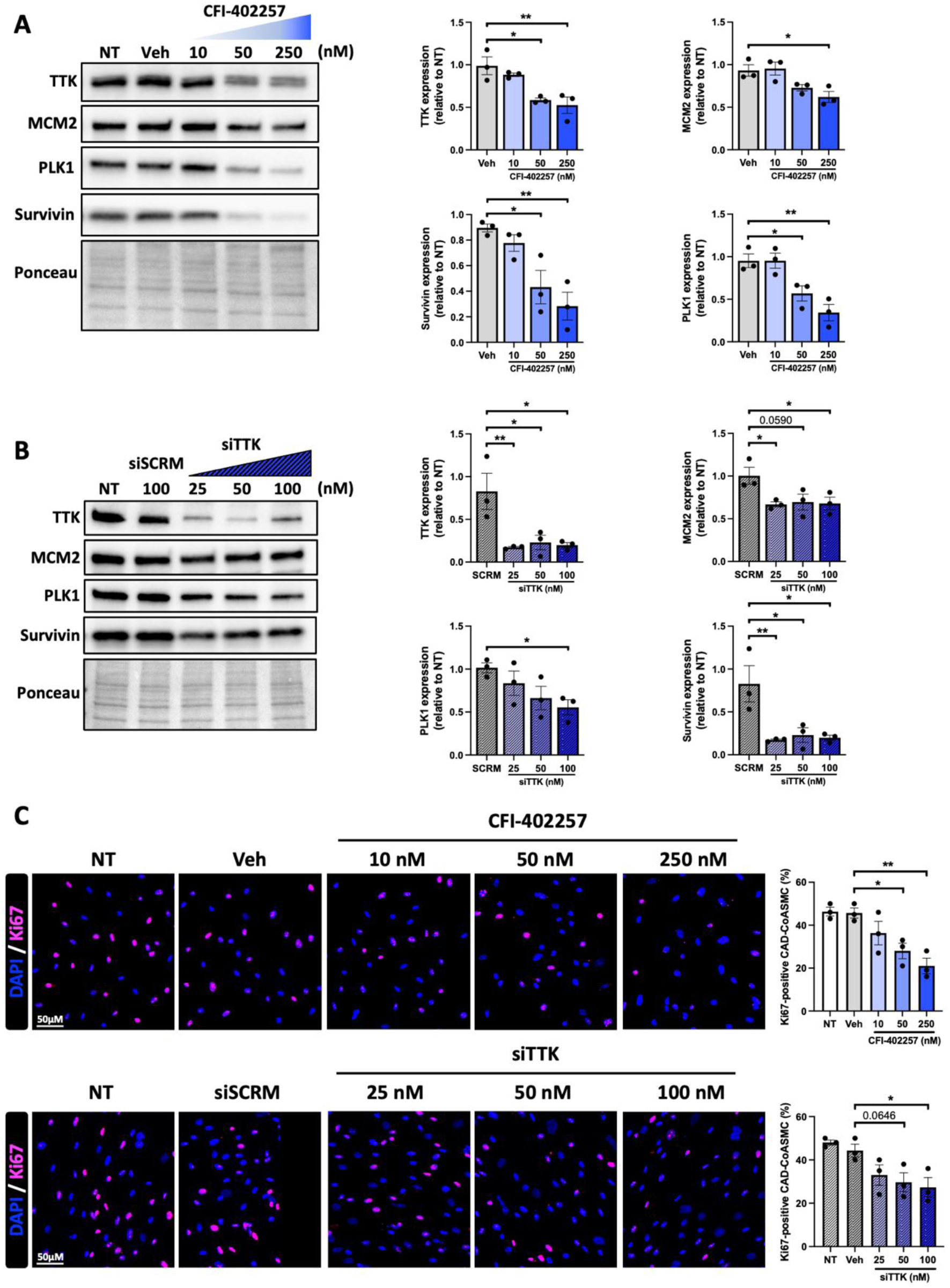
Pharmacological and molecular inhibition of TTK decrease pro-proliferative phenotype of CAD-CoASMCs. **(A-B)** Western blot and corresponding quantification of TTK, MCM2, PLK1 and survivin in CAD-CoASMCs exposed for 48h to escalating concentration of (A) CFI-402257 (TTK inhibitor) and (B) siTTK *(n=3, *P < 0.05, **P < 0.0, one-way ANOVA followed by Tukey’s post hoc analysis; data represent mean ± SEM)*. **(C)** Representative fluorescent images and corresponding quantification of Ki67- labeled (red) of CAD-CoASMCs exposed or not exposed to escalating concentration of CFI-402257 and siTTK for 48h. Cell nuclei were counterstained with DAPI (blue) (*n = 3; *P < 0.05, **P < 0.01, one-way ANOVA followed by Tukey’s post hoc analysis; data represent mean ± SEM*). Scale bar, 50μm

Together, these findings establish that TTK is a key downstream effector of hypusine-dependent translational control, and that inhibition of DHPS suppresses the CAD-CoASMC proliferative program in part through TTK downregulation, thereby recapitulating the antiproliferative effects achieved by direct TTK inhibition.

### Pharmacological inhibition of DHPS using GC7 attenuates carotid stenosis in vivo

To evaluate the therapeutic potential of hypusine pathway inhibition *in vivo*, we assessed the effects of DHPS blockade on vascular remodeling following carotid injury. Adult rats underwent right carotid endothelial denudation using a flexible wire to induce robust neointimal hyperplasia, then received daily injections of GC7 or vehicle for two weeks (Figure 5A). As expected, vehicle-treated rats exhibited pronounced neointimal hyperplasia compared with sham controls, as visualized by elastica Van Gieson (EVG) staining (Figure 5B). In contrast, GC7 treatment markedly prevented neointimal hyperplasia, reflected by a smaller neointima area without any change in the media area, a decrease of the ratio neointima-to-media and an improvement of the luminal obliteration (Figure 5B–C). Toxicity assessments revealed no significant differences in kidney or liver function between GC7- and vehicle-treated animals (Figure S6). These results are in agreement with a previous work showing no adverse effects on cardiac structure or systemic health following GC7 treatment^18^. Together, these findings demonstrate that pharmacological DHPS inhibition with GC7 effectively mitigates neointimal hyperplasia without detectable toxicity, supporting its potential as a therapeutic strategy for restenosis and other vascular remodeling disorders.

**Figure 5.**
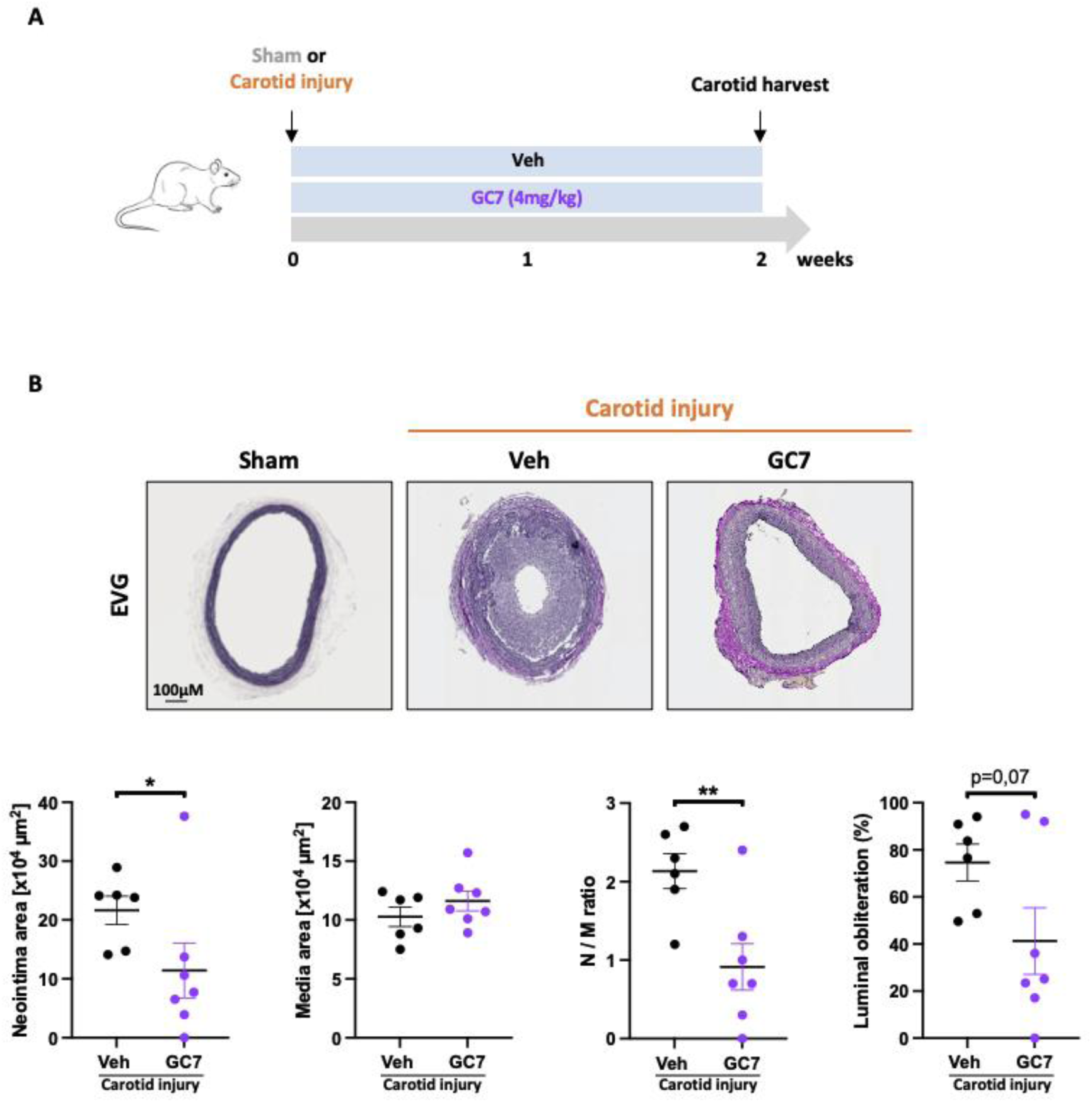
Pharmacological inhibition of DHPS improves vascular remodeling induced by carotid injury. **(A)** Schematic diagram of the carotid artery injury experiment in Sprague-Dawley rats. **(B)** Representative images of carotid arteries stained with elastica van gieson (EVG) in rats subjected or not to carotid injury and treated or not with GC7 (4mg/kg) for 2 weeks and corresponding quantification of neointima area, media area, N/M ratio and luminal obliteration (*n = 6 to 7; *P < 0.05 and **P < 0.01, unpaired Student’s t-test; data represent mean ± SEM*). Scale bar, 100μm.

### *Dhps* loss-of-function targeted to smooth muscle cells alleviates carotid stenosis in mice

To assess whether DHPS deletion specifically in VSMCs is required for neointimal hyperplasia, we generated SMC targeted *Dhps*-deficient heterozygous mice (*Dhps^flox/+^;Tg^+/Tagln-Cre^*), since homozygous SMC-specific *Dhps* deletion is not viable^18^. These mice were phenotypically indistinguishable from their wild-type (WT) counterparts (*Dhps^flox/flox^, Dhps^flox/+^, or Tg^+/Tagln-Cre^).* All animals underwent total carotid ligation and were sacrificed 28 days post-injury, the time point previously shown to yield complete vessel obliteration^9^ (Figure 6A). Immunostaining confirmed a decrease in DHPS expression in *Dhps^flox/+^;Tg^+/Tagln-Cre^* carotids (Figure 6B). Histomorphometric analysis revealed that SMC-specific *Dhps* loss significantly prevented neointimal hyperplasia, as demonstrated by a decrease of neointima area, the neointima/media ratio and luminal obliteration, without affecting media area (Figure 6C).

**Figure 6.**
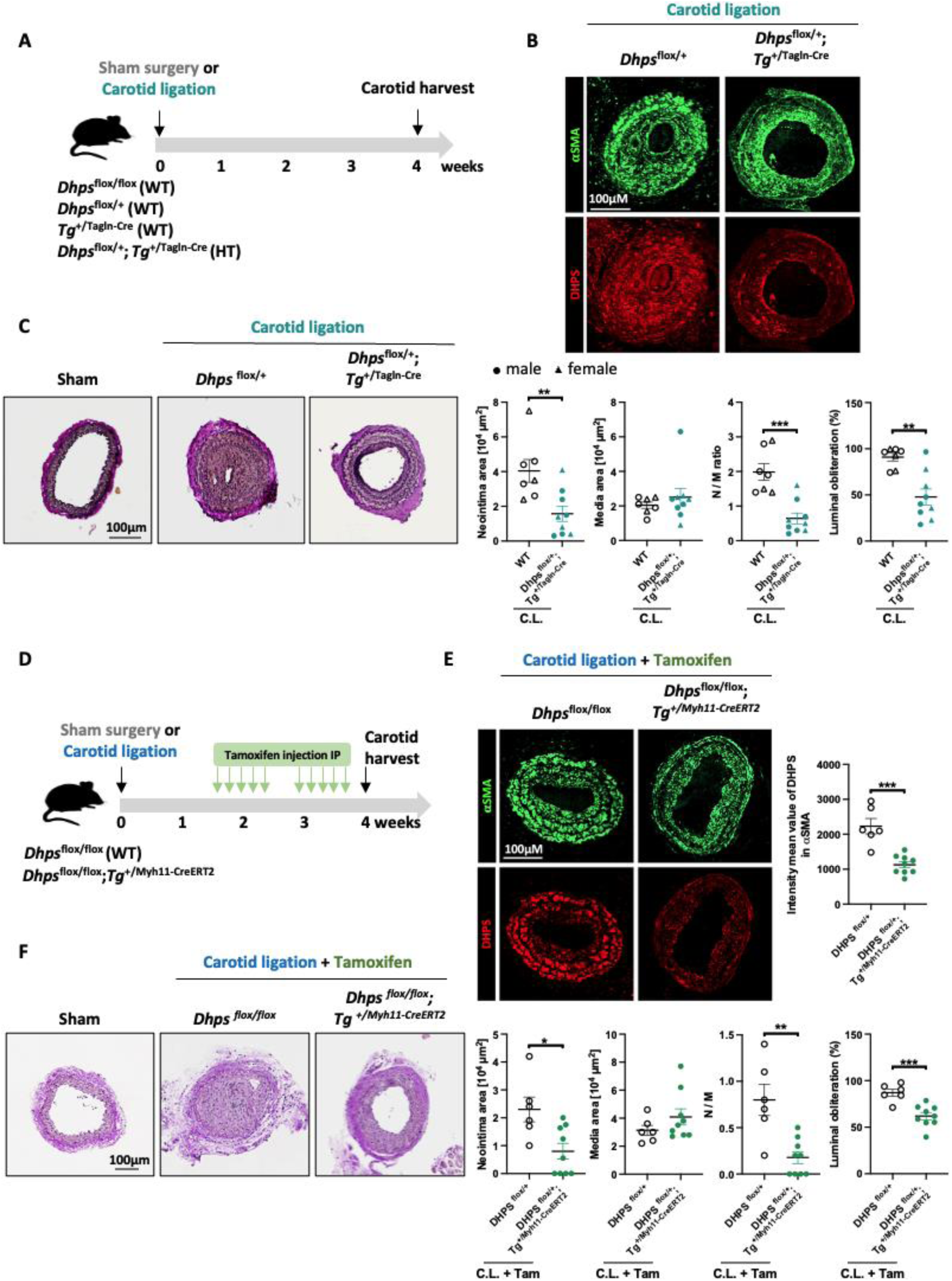
*Dhps* loss of function targeted to smooth muscle cells improve vascular remodeling induced by carotid ligation. **(A)** Schematic of the carotid ligation protocol in WT (*Dhps^flox/flox^*, *Dhps^flox/+^* and *Tg^+/Tagln-Cre^*) and *Dhps^flox/+^;Tg^+/Tagln-Cre^*male and female mice. **(B)** Immunofluorescence images of left carotid arteries labelled for DHPS (red) and αSMA (green) in *Dhps^flox/+^*(WT) and *Dhps^flox/+^;Tg^+/Tagln-Cre^*mice subjected to carotid ligation. Scale bar, 100µm. **(C)** Representative images of the carotid arteries stained with Elastica Van Gieson (EVG) in WT and *Dhps^flox/+^;Tg^+/Tagln-Cre^*mice subjected or not to carotid ligation at week 4 and corresponding quantification of neointima area, media area, N/M ratio and luminal obliteration (*n = 7 to 9; **P < 0.01 and ***P < 0.001, unpaired Student’s t-test; data represent mean ± SEM*). Scale bar, 100μm. **(D)** Schematic of the carotid ligation protocol in WT (*Dhps^flox/flox^*) and *Dhps^flox/flox^;Tg^+/Myh11-creERT2^* male mice. The first tamoxifen injection to induce Dhps deletion specifically in smooth muscle cells was administered on day 10 post-injury. A total of ten tamoxifen injections were realised, in two rounds of five consecutive daily injections, separated by a two-day rest period. **(E)** Representative immunofluorescence images of left carotid arteries labelled for DHPS (red) and αSMA (green) in tamoxifen-treated WT (*Dhps^flox/flox^*) and *Dhps^flox/flox^;Tg^+/Myh11-creERT2^*male mice subjected to carotid ligation and corresponding quantification of DHPS intensity in αSMA (*n = 6 to 9; ***P < 0.001, unpaired Student’s t-test; data represent mean ± SEM*) Scale bar, 100µm. **(F)** Representative images of the carotid arteries stained with EVG in tamoxifen-treated WT (*Dhps^flox/flox^*) and *Dhps^flox/flox^;Tg^+/Myh11-creERT2^*male mice subjected or not to carotid ligation at week 4 and corresponding quantification of neointima area, media area, N/M ratio and luminal obliteration (*n = 6 to 9;*P < 0.05, **P < 0.01 and ***P < 0.001, unpaired Student’s t-test; data represent mean ± SEM*). Scale bar, 100μm.

As a complementary approach, we next evaluated whether inducible deletion of *Dhps* in adult SMCs could prevent disease progression after injury. *Dhps^flox/flox^* mice were crossed with Myh11-CreERT2 mice to generate *Dhps^flox/flox^;Tg^+/Myh11-CreERT2^* animals. Carotid ligation was performed, and tamoxifen was administered starting on day 10, after the onset of neointimal hyperplasia^9^, and animals were sacrificed on day 28 (Figure 6D). Efficient deletion of *Dhps* was confirmed by immunostaining (Figure 6E). As in the heterozygous mice, inducible *Dhps* deletion significantly improved neointimal hyperplasia (reduction of neointima area, the neointima/media ratio, and luminal obstruction, Figure 6F).

Collectively, these results establish that DHPS expression in VSMCs contributes to the development and progression of neointimal hyperplasia and underscore DHPS as a promising therapeutic target for restenosis.

### DHPS inhibition prevents vascular remodeling in cultured human coronary artery and saphenous vein rings

To examine the translational relevance of hypusine pathway inhibition, we used *ex vivo* cultures of human CoA and SV rings, which preserve native multicellular architecture and allow the assessment of drug effects over several days. In these experiments, CoA segments were obtained from hearts of CAD patients undergoing transplantation. CoAs rings were mechanically injured and stimulated with PDGF-BB and FGF2 to induce pathological inward remodeling^9^. As expected, neointimal hyperplasia was increased, evidenced by larger neointima area without changes in media area, resulting in an increased neointima-to-media ratio and luminal narrowing (Figure 7A). This remodeling was accompanied by elevated hypusinated eIF5A and proliferation markers (Figure 7B). Co-treatment with GC7 effectively attenuated these changes, reducing neointima area, the ratio neointima-to-media, and resulting in luminal preservation. The structural improvements were associated with reduced eIF5A hypusination, increased eIF5A acetylation, and decreased expression of proliferation and survival markers (Figure 7B). We next evaluated human SV rings obtained during CABG. Tissues were mechanically injured and cultured with a growth factor cocktail containing PDGF-BB, FGF2, and 7-ketocholesterol. Since internal elastin is challenging to visualize, we limited our assessment to measuring wall thickness and luminal narrowing. As anticipated, stimulation induced vessel wall thickening and luminal narrowing (Figure 8A), alongside increased hypusinated eIF5A and expression levels of proliferation proteins (Figure 8B). GC7 co-treatment significantly blunted these molecular and structural alterations, reducing both vascular remodeling and expression of pathological markers (Figure 8A and B).

**Figure 7.**
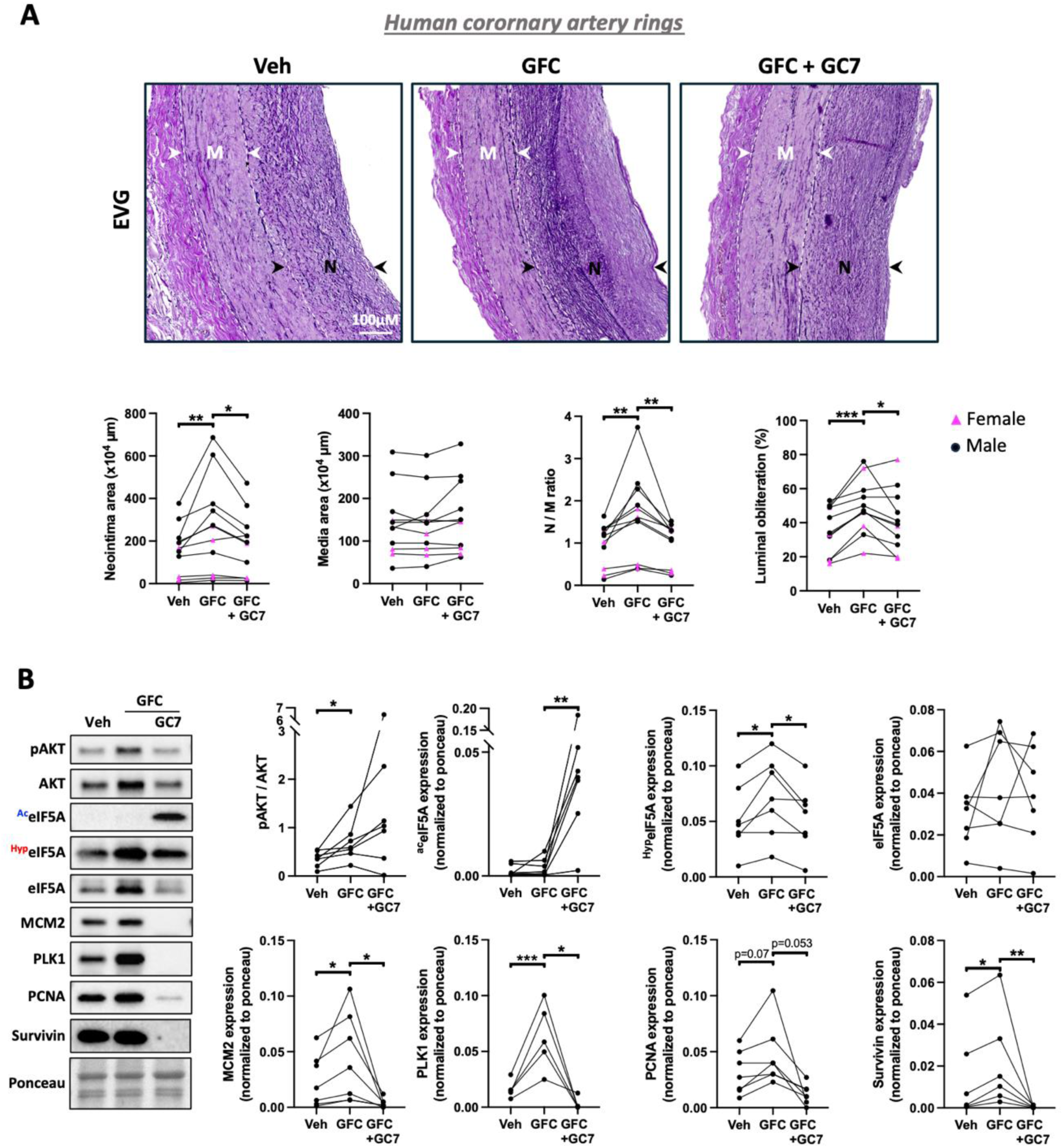
Pharmacological inhibition of DHPS reduces neointimal hyperplasia in cultured human coronary artery rings. **(A)** Representative images of human coronary artery rings stained with Elastica Van Gieson (EVG) after exposure to GFC (PDGF-BB and FGF2, both at 30ng/mL) in presence or absence of GC7 (30µM) for 5 days, and corresponding quantification of neointima area, media area, N/M ratio and luminal obliteration. The external and internal elastin are delineated with dashed lines. The white arrows indicate the media area (M), while the black arrows mark the neointima area (N) (n = 11). Scale bar, 100μm. **(B)** Representative western blots et corresponding quantification of pAKT, AKT, ^Ac^eIF5A, ^Hyp^eIF5A, eIF5A, MCM2, PLK1, PCNA and survivin in human coronary artery rings exposed to GFC (PDGF-BB and FGF2; both at 30ng/mL) in presence or absence of GC7 (30µM) for 5 days (n= 5 to 7). The normality hypothesis was verified using the Shapiro-Wilks test using residuals from the statistical model and transformed by the Cholesky’s metric. The Brown and Forsythe’s variation of Levene’s test statistic was used to verify the homogeneity of variances. pAKT/AKT, ^Ac^EIF5A, MCM2, PLK1 and survivin were log-transformed to fulfill the normality and variance assumptions. Statistical significance was declared significant with a two-tailed *p* value < 0.05 (*\*P < 0.05, **P < 0.01, ***P < 0.01, data represent mean ± SEM)*

**Figure 8.**
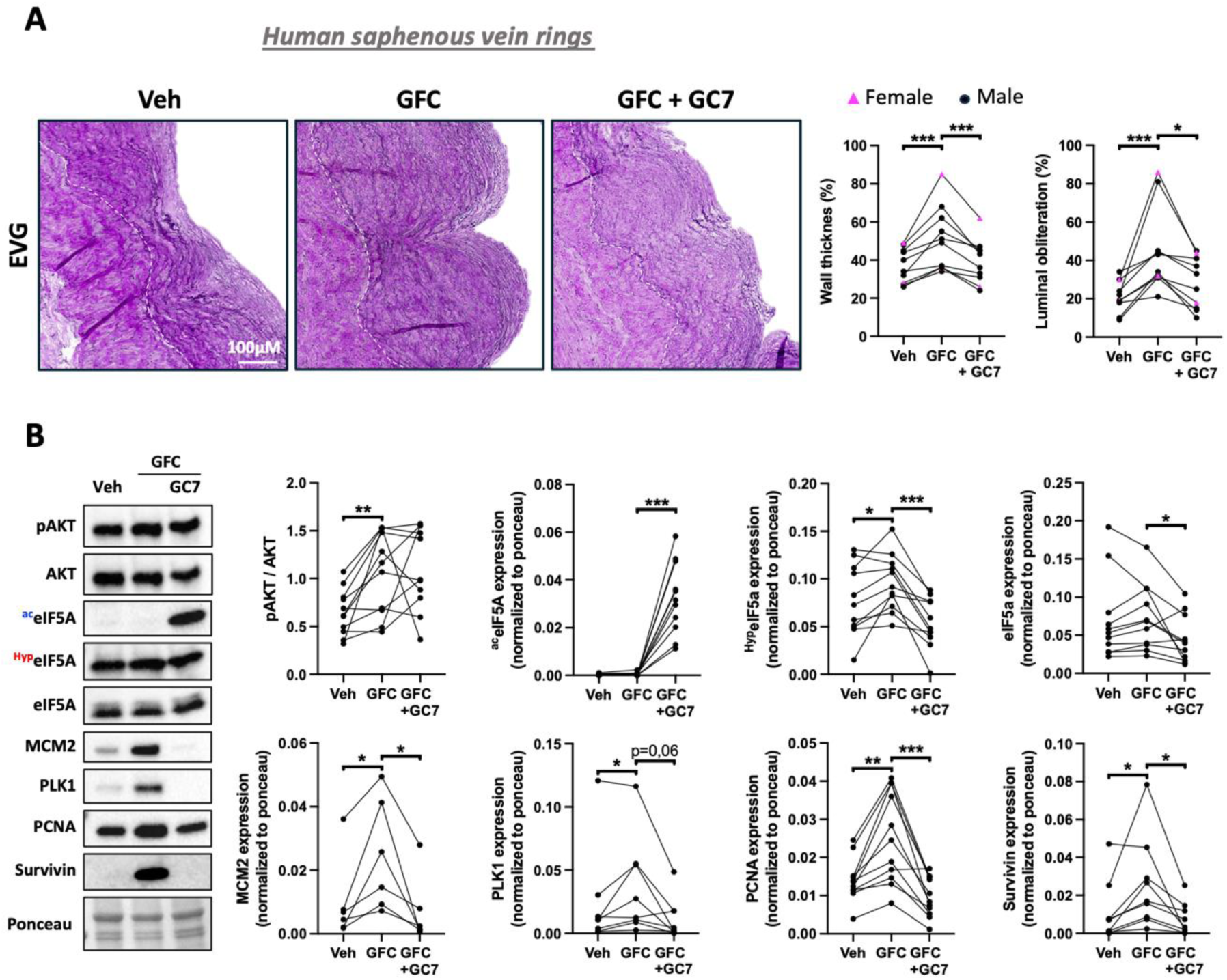
Pharmacological inhibition of DHPS reduces vascular remodeling in cultured human saphenous vein rings. **(A)** Representative images of human saphenous vein rings stained with elastica van gieson (EVG) after exposure to GFC (PDGF-BB and FGF2, both at 30ng/mL + 7-Ketocholesterol 10µM) in presence or absence of GC7 (30µM) for 5 days, and corresponding quantification of wall thickness and luminal obliteration. The external elastin is delineated with white dashed lines (n = 10). Scale bar, 100μm. **(B)** Representative western blots et corresponding quantification of pAKT, AKT, ^Ac^eIF5A, ^Hyp^eIF5A, eIF5A, MCM2, PLK1, PCNA and survivin in human saphenous vein rings exposed to GFC (PDGF-BB and FGF2, both at 30ng/mL + 7-Ketocholesterol 10µM) in presence or absence of GC7 (30µM) for 5 days (n= 6 to 11). The normality hypothesis was verified using the Shapiro-Wilks test using residuals from the statistical model and transformed by the Cholesky’s metric. The Brown and Forsythe’s variation of Levene’s test statistic was used to verify the homogeneity of variances. MCM2, PLK1 and survivin were log-transformed to fulfill the normality and variance assumptions. Statistical significance was declared significant with a two-tailed *p* value < 0.05 (*\*P < 0.05, **P < 0.01, ***P < 0.01, data represent mean ± SEM)*

Together, these results demonstrate that human *ex vivo* vessel culture faithfully model pathological remodeling and that GC7 inhibits hypusine signaling, suppressing both structural and molecular features of disease progression. These findings provide strong translational support for DHPS inhibition as a therapeutic approach in human vascular remodeling disorders.

## DISCUSSION

Excessive proliferation of VSMCs is a key driver of restenosis and other occlusive vascular diseases. Despite significant progress in endovascular therapy, restenosis remains a major clinical limitation, largely due to maladaptive VSMC responses to vascular injury. Identifying molecular pathways and key driver that govern VSMC phenotypic changes toward pro-proliferative state is therefore essential for developing next-generation therapies. In this study, we demonstrate that hypusine signaling is markedly upregulated in human CAD-CoASMCs and is associated with enhanced proliferation and sustained ECM production. Inhibition of this pathway through pharmacological targeting of DHPS with GC7 mitigated the proliferative phenotype and significantly attenuated neointimal hyperplasia in both *in vivo* and *ex vivo* models of restenosis. These findings position the hypusine pathway as a compelling therapeutic target for mitigating restenosis in CAD.

Our results align with prior work by Lemay et al.^18^, who reported that inhibition of eIF5A hypusination exerts antiproliferative effects in pulmonary artery SMCs and improves pulmonary vascular remodeling in experimental pulmonary arterial hypertension. Because hypusination depends on spermidine from the polyamine biosynthetic pathway, our findings also intersect with prior studies showing that blockade of polyamine production using DFMO, an inhibitor of ODC1 upstream of spermidine synthesis, reduces VSMC proliferation and vascular remodeling in murine restenosis models^19–21^ (Figure S1). Our results suggest that the therapeutic benefits of polyamine pathway inhibition may, at least in part, be mediated through suppression of eIF5A hypusination. Targeting hypusination directly may therefore produce similar antirestenotic effects while avoiding broader physiological disruption associated with global suppression of polyamine metabolism.

Inflammation is another key determinant of vascular remodeling in atherosclerosis and restenosis, with macrophages promoting VSMC proliferation and migration^42–45^. Hypusine signaling has known roles in immune regulation: DHPS and hypusinated eIF5A enhance inflammatory macrophage phenotypes in obesity^46,47^ and promote M2 polarization in tumor-associated macrophages to sustain tumor growth^48,49^. Thus, inhibition of hypusination in macrophages could theoretically contribute to the improvements observed in our *in vivo* models. However, our *ex vivo* human vascular tissue experiments, performed in the absence of inflammatory cells, still demonstrated reduced neointimal development and decreased markers of proliferation, indicating that the primary therapeutic effects of hypusination blockade arise from direct modulation of VSMCs.

An additional observation from our study is the marked increase in eIF5A acetylation when hypusination is inhibited. Hypusination promotes the cytoplasmic localization of eIF5A and its translation activity^50,51^, whereas acetylation drives its nuclear translocation^51^ and can, in some contexts, lead to degradation^52^. Despite enhanced acetylation, we did not observe a reduction in total eIF5A protein levels in our models, suggesting nuclear accumulation rather than degradation. The functional role of eIF5A in the nucleus remains poorly understood and represents an important area for future investigation.

This study also has limitations. We did not dissect the individual contributions of the two eIF5A isoforms: eIF5A1 and eIF5A2, are overexpressed in several pathological conditions^49,53–55^, open the possibility of isoform-specific effects. Although GC7 is a widely used DHPS inhibitor, it is not fully selective, and off-target effects cannot be entirely excluded. Moreover, our *ex vivo* human tissue models lack immune components that contribute to restenosis, particularly macrophages. The number of primary CoASMC lines used for proteomic studies was relatively limited, which may not fully represent the heterogeneity of CAD. Finally, long-term safety, pharmacokinetics, and metabolic consequences of sustained DHPS inhibition were not assessed, underscoring the need for further preclinical evaluation prior to clinical translation.

In conclusion, this study identifies hypusine signaling as a central regulator of the re-oriented pro-proliferative phenotype of VSMC and restenosis in CAD, providing strong proof-of-concept that DHPS inhibition can effectively prevent restenosis. The reproducibility of these findings across cellular, animal, and human tissue models highlights the translational potential of targeting this pathway. Future work should focus on dissecting isoform-specific functions of eIF5A, elucidating the nuclear role of acetylated eIF5A, and developing DHPS inhibitors with high specificity and favorable pharmacokinetic profiles suitable for clinical use.

## Acknowledgements

We thank the IUCPQ Biobank of the Quebec Respiratory Health Research Network as well as the department of cytology and pathology from the IUCPQ for providing access to tissue and clinical data.

## Abbreviations

ACTG2: Smooth muscle actin gamma-2
CABG: Coronary artery bypass grafting
CAD: Coronary artery disease
CDK2: Cyclin-dependent kinase 2
CoA: Coronary artery
CoASMC: Coronary artery smooth muscle cell
DHPS: Deoxyhypusine synthase
DOHH: Deoxyhypusine hydroxylase
ECM: Extracellular matrix
eIF5A: Eukaryotic translation initiation factor 5A
EVG: Elastica Van Gieson
GC7: N1-guanyl-1,7-diaminoheptane
MCM2: Minichromosome maintenance complex component 2
PLK1: Polo-like kinase 1
PCNA: Proliferating cell nuclear
SV: Saphenous vein
TTK: Threonine tyrosine kinase
UHRF1: Ubiquitin-like containing PHD and RING finger domains 1
VSMC: Vascular smooth muscle cell

